# A NOVEL MACHINE LEARNING APPROACH FOR TUMOR DETECTION BASED ON TELOMERIC SIGNATURES

**DOI:** 10.1101/2025.05.28.656542

**Authors:** Priyanshi Shah, Arun Sethuraman

**Affiliations:** Department of Biology, San Diego State University, San Diego, California, 92182, United States of America

## Abstract

Cancer remains one of the most complex diseases faced by humanity, with over 200 distinct types, each characterized by unique molecular profiles that demand specialized therapeutic approaches (Tomczak et al., 2015). Prior studies have shown that both short and long telomere lengths are associated with elevated cancer risk, underscoring the intricate relationship between telomere length variation and tumorigenesis (The Telomeres Mendelian Randomization Collaboration et al., 2017). To investigate this relationship, we developed a supervised machine learning model trained on telomeric read content, genomic variants, and phenotypic features to predict tumor status. Using data from 33 cancer types within The Cancer Genome Atlas (TCGA) program, our model achieved an accuracy of 82.62% in predicting tumor status. The trained model is available for public use and further development through the project’s GitHub repository: https://github.com/paribytes/TeloQuest. This work represents a novel, multidisciplinary approach to improving cancer diagnostics and risk assessment by integrating telomere biology with Biobank-scale genomic and phenotypic data. Furthermore, we highlight the potential of telomere length variation as a meaningful predictive biomarker in oncology.

## INTRODUCTION

In middle- and high-income countries, approximately one in three people will develop clinical cancer during their lifetime (Greaves, 2007). This high occurrence of cancer in modern society presents a complex interplay of genetics and elevated mutation load due to rapid environmental shifts caused by factors such as pollution, and lifestyle changes. (Aviv et al., 2017). Nonetheless, cancers have strong heritable, biotic contributors, including telomeric length variation (Victorelli & Passos, 2017). Telomeres in vertebrates consist of tandem hexameric sequence repeats (TTAGGG)_n_ located at the ends of linear chromosomes (Victorelli & Passos, 2017). These repetitive nucleotide sequences safeguard chromosome ends and uphold genomic stability (Aunan et al., 2016). In human somatic cells, telomeres gradually shorten with each cell division due to DNA replication, ultimately triggering cellular senescence or programmed cell death once a critical length is reached (Okamoto & Seimiya, 2019). This process underpins an evolutionary trade-off, where protection against cancer is achieved at the expense of reduced regenerative capacity (Gomes et al., 2011). In contrast, self-renewing cells such as embryonic stem cells, and most notably, cancer cells can overcome telomere shortening and maintain cellular immortality by sustaining telomere length, typically through the upregulation of telomerase (Okamoto & Seimiya, 2019; Shay & Wright, 2011). Replicative telomere attrition thus serves as a critical tumor-suppressive mechanism in normal cells, acting as a barrier to unchecked proliferation (Xu et al., 2013).

However, when this protective mechanism fails, short telomeres can become a source of genomic instability, contributing to cancer development (Aviv et al., 2017). Individuals with short telomeres have been shown to be at increased risk of various cancers, including gastrointestinal tumors (Kroupa et al., 2019), bladder cancer (McGrath et al., 2007) and head and neck cancer (Zhu et al., 2016). Interestingly, studies have identified a significant inverse association between short telomeres and lung cancer risk, suggesting that telomere-related carcinogenic mechanisms may vary across different cancer types (Zhu et al., 2016). Additionally, short telomeres are implicated in a broader disease phenotype affecting individuals from infancy to adulthood, with more severe outcomes observed in younger patients (Stanley & Armanios, 2015). Patients with short telomeres may also exhibit immunosenescence, impairing cancer surveillance and further increasing susceptibility to malignancies (Stanley & Armanios, 2015). These findings support the widely held view that genomic instability arising from critically short telomeres, especially when combined with other oncogenic factors plays a central role in cancer development (Aviv et al., 2017).

On the other hand, telomere shortening may function as a tumor-suppressive mechanism by limiting cellular proliferation, while individuals with longer telomeres may face a higher risk of acquiring somatic mutations due to increased proliferative capacity (Maciejowski & De Lange, 2017; The Telomeres Mendelian Randomization Collaboration et al., 2017). For instance, several studies have shown that individuals with longer telomeres are more likely to develop melanoma, suggesting that shorter telomeres may offer protection against malignant transformation by promoting cellular senescence (Rachakonda et al., 2018; Zhu et al., 2016). Similarly, long telomeres have been associated with an elevated cancer risk in vertebrate animal models (McNally et al., 2019). These findings support the biological plausibility that telomeres play a two-edged sword in cancer development, contributing to a paradoxical relationship between telomere length and cancer risk (Zhu et al., 2016).

Telomere length (TL) is itself a complex and highly heritable genetic trait, with significant variation across modern human populations (Factor-Litvak et al., 2016). Evolutionary forces may have contributed to these differences across populations; for example, polygenic adaptation may explain the shorter TL observed in individuals of European ancestry compared to those of African ancestry, potentially as a protective adaptation against the higher risk of melanoma in depigmented skin types (Hansen et al., 2016; Mangino et al., 2015). Furthermore, McQuillan et al. (2024) reported that adults from indigenous populations in malaria-endemic regions of Africa had shorter TL than those from malaria-free areas, suggesting environmental selection pressures on telomere dynamics.

TL also plays a significant role in regulating cellular proliferative capacity, mutation accumulation, and cancer susceptibility (Aviv et al., 2017). Irregularities in TL are increasingly recognized as contributors to clinically significant disease processes (McNally et al., 2019). However, the consistent exclusion of TL from mathematical models of mutation risk may reflect challenges in distinguishing between constitutive TL – measured in normal somatic tissues – and TL in tumor cells (Aviv et al., 2017). Precise phenotypic characterization of TL is therefore essential for advancing precision medicine, essentially in oncology. Despite this, much of the epidemiological data on TL in cancer relies on quantitative polymerase chain reaction (qPCR), a method with known variability across laboratories, potentially limiting reproducibility and cross-study comparisons (Verhulst et al., 2015).

According to Aviv et al. (2017), simply knowing whether TL is inherently long or short in individuals with cancer is not sufficient for understanding the role of telomeres in the disease. Additionally, it’s crucial to have precise and reproducible metrics for TL to understand the absolute differences in TL between cancer cases and controls, as well as between cancerous and non-cancerous tissues within individuals. This information is important to gain insights into cancer progression and the potential role of telomeres in its regulation. Besides cancer-specific correlations, TL may influence cancer risk in a multifaceted manner rather than through a straightforward linear relationship (Zhu et al., 2016).

Here, we present a supervised machine learning approach to explore the complex, and sometimes contradictory relationships between telomere length dynamics and human cancers. Telomere content (or telomeric read content, the scaled number of sequencing reads mapped back to telomeric regions), used as a proxy for telomere length was estimated using qmotif v1.0 (Holmes et al., 2022) by analyzing whole genome sequencing (WGS) data from The Cancer Genome Atlas (TCGA) program. The TCGA is a comprehensive resource containing genomic and clinical data from more than 30 types of human tumors (Tomczak et al., 2015). It has enabled both cancer-specific and pan-cancer analyses, significantly advancing our understanding of tumor biology and helping drive improvements in how cancer is diagnosed, treated, and prevented (Tomczak et al., 2015). In total, we analyzed telomere content across 17,400 unique samples derived from 8,816 patients representing 33 cancer types and apply this in combination with genomic and phenotypic variables to build a predictive model of tumor status.

## METHODS

### Genomic Data Access, BAM Slicing, Estimating Telomere Content

We utilized the TCGA “Cohort Builder” tool within the Genomic Data Commons (GDC) Data Portal to access genomic, phenotypic, and diagnostic information for each project (categorized by cancer type). Chromosomal coordinates were converted from GRCh37 to GRCh38 using UCSC’s LiftOver (Hinrichs, 2006) to be congruent with the human telomeric coordinates specified in the qmotif v1.0 configuration file. Next, the GDC’s ‘BAM Slicing API’ was employed to extract genomic data relevant to the converted telomeric regions across all cohorts, only pertinent to the “lifted over” telomeric coordinates. Using samtools v1.12 (Danecek et al., 2021), we generated BAI index files for all the sliced BAM files. Finally, we applied the qmotif pipeline (Supplementary Figure S1) to sequentially process the BAM and BAI files, estimating scaled telomeric reads - defined as the normalized number of sequencing reads mapping to telomeric regions.

### Extracting Variants across 15 Telomere Length Related Genes, Genotype Encoding

We also extracted genomic single nucleotide variants from BAM files at 15 genes associated with telomere shortening and elongation, as identified in a recent study (Burren et al., 2024; Table 2). Chromosomal coordinates for these genes were retrieved using the UCSC’s Genome Browser, based on GRCh38 reference genome build (Supplementary Table S2). Variant calling was then performed using bcftools v1.21 (Danecek et al., 2021), applying the mpileup command under the assumption of diploidy at all sites. To assure consistency across analyses, the GRCh38 reference genome FASTA file was sourced from the GDC Reference Files repository (https://gdc.cancer.gov/about-data/gdc-data-processing/gdc-reference-files). Genotype encoding was subsequently applied using a standardized numeric code as described in Table 1.

**Table 1.** Genotype Encoding.

| Genotype notation | Name | Numeric code | Description | Importance |
| --- | --- | --- | --- | --- |
| 0/0 | Homozygous reference | 0 | Indicates two copies of the reference allele at a specific genomic location. | Considered the “common” or major allele across the population. |
| 0/1 | Heterozygous | 1 | Indicates one reference allele and one alternate allele. It signifies genetic variation, as it includes a mutation or variation from the reference. | Heterozygosity is biologically important because it can contribute to genetic diversity (Hedrick, 2006) and, depending on the variant, might affect traits or susceptibility to diseases (Kidd et al., 2012). |
| 1/1 | Homozygous alternate | 2 | Indicates both alleles at this locus are the alternate (variant) allele. | Can be biologically significant if the alternate allele is associated with traits or diseases. Homozygous alternate genotypes may have stronger or different effects than heterozygous genotypes, especially if the variant has a recessive or additive effect (Stranger et al., 2012). |
| ./. | Missing data | None | Represents missing values. | - |

**Table 2.** Gene symbol, name, and summary of the functions of the 15 telomeric length related genes for which variants have been captured from the TCGA dataset. The information was collated using https://www.ncbi.nlm.nih.gov/gene/.

| Gene Symbol | Gene Full Name | Function |
| --- | --- | --- |
| DCLRE1B | DNA cross-link repair 1B | Involved in repair of interstrand cross-links |
| NAF1 | Nuclear assembly factor 1 ribonucleoprotein | Involved in telomerase RNA stabilization |
| TERT | Telomerase reverse transcriptase | Maintenance of telomere ends |
| G3BP1 | G3BP stress granule assembly factor 1 | ATP-dependent DNA/RNA helicase in Ras signaling pathway. |
| ZNF451 | Zinc finger protein 451 | Enables SUMO ligase activity |
| POT1 | Protection of telomeres 1 | Regulates telomere length and protects chromosome ends |
| TERF1 | Telomeric repeat binding factor 1 | Telomere protein inhibiting telomerase, regulates chromosome ends. |
| STN1 | STN1 subunit of CST complex | Functions in telomere-associated complexes and DNA replication |
| ATM | ATM serine/threonine kinase | Master controller of cell cycle checkpoints |
| TINF2 | TERF1 interacting nuclear factor 2 | Protection of telomeres |
| PARN | Poly(A)-specific ribonuclease | Telomere maintenance, gene expression, disease risk |
| CTC1 | CST telomere replication complex component 1 | Protects telomeres from degradation |
| BRIP1 | BRCA1 interacting DNA helicase 1 | Normal double-strand break repair function of breast cancer |
| SAMHD1 | SAM and HD domain containing deoxynucleoside triphosphate triphosphohydrolase 1 | Mediation in tumor necrosis |
| RTEL1 | Regulator of telomere elongation helicase 1 | Protection and elongation of telomeres |

To assess mutational load and reduce dimensionality of features used in our machine learning models, we calculated three summary statistics for each sample: total heterozygous mutations, total homozygous alternate mutations, and total mutations (the sum of homozygous and heterozygous mutations).

Telomeric read content, genomic variant summaries, and phenotypic and diagnostic data were then consolidated into a single CSV file. The phenotypic and diagnostic features used for model training included age at diagnosis, ethnicity, gender, race, vital status, primary diagnosis, prior malignancy, prior treatment, site of resection or biopsy, synchronous malignancy, tissue or organ of origin, treatment, treatment type, and tumor status (see Supplementary Table S1 for the full list of variables).

### Model Training and Testing

A supervised machine learning approach was employed to predict tumor status across all 33 TCGA cancer types using a Random Forest classifier (additional models tested included a logistic regression-based classifier, and a support vector machine-based classifier, which either under- or over-fit data – see Supplement). Our model was trained and tested on the compiled dataset of 17,400 samples in the CSV file (Supplementary Table S3).

### Data Preparation and Imputation

The dataset was split into training and test sets using an 80/20 ratio. To address missing data, numerical features, including telomere content, total homozygous alternate mutations, total heterozygous mutations, and total mutations were imputed using mean values. Categorical features such as ethnicity, race, tissue or organ of origin, and primary diagnosis were imputed using the most frequent category within each respective feature.

### Hyperparameter Tuning and Model Optimization

Hyperparameter tuning was conducted to enhance the model’s performance across evaluation metrics such as accuracy, precision, recall, and F-1 score. We used the GridSearchCV tool from the scikit-learn library, which automates the search for the best combination of hyperparameters through cross-validation. Tuned parameters included: n_estimators (number of trees in the forest), max_depth (maximum depth of each tree), max_features (number of features considered at each split), min_samples_split (minimum number of samples required to split a node), oob_score = True (enables out-of-bag score estimation for model accuracy), and bootstrap = True (enables sampling with replacement to increase tree diversity and model robustness). To evaluate model generalizability, we applied 10-fold cross validation, dividing the dataset into 10 equal subsets, with each fold used once as a test set while the remaining folds were used for training. The Random Forest model was implemented using the RandomForestClassifier in scikit-learn in a Jupyter notebook environment.

### Threshold Tuning and Model Evaluation

Threshold tuning involves adjusting the classification threshold used to convert predicted probabilities into class labels. While a default threshold of 0.5 is commonly used, classifying probabilities ≥0.5 as positive and <0.5 as negative, this objective cutoff may not be ideal in all scenarios. Particularly when the cost of false positives and false negatives differ, or when specific performance metrics (e.g., sensitivity, specificity, or F1 score) are prioritized, a customized threshold can yield improved results. To optimize our model’s classification performance, we evaluated a range of thresholds using the best-performing hyperparameter combination identified via GridSearchCV as described above. Model performance at each threshold was then assessed using accuracy, precision, recall (sensitivity), specificity, and F1 score. Additionally, an ROC curve was generated to visualize model performance across all thresholds (Figure 3).

To determine the optimal threshold, we computed Youden’s Index, a metric that maximizes the sum of sensitivity (true positive rate) and specificity (true negative rate). This approach identifies the threshold that achieves the best trade-off between correctly identifying positive cases (sensitivity) and minimizing false positives (specificity). Finally, to assess the contribution of each input feature variable to the model’s predictions, we generated a feature importance plot (Figure 2). This visualization ranks all features in the model in descending order of their influence, offering insight into the most impactful predictors of tumor status.

**Figure 1.**
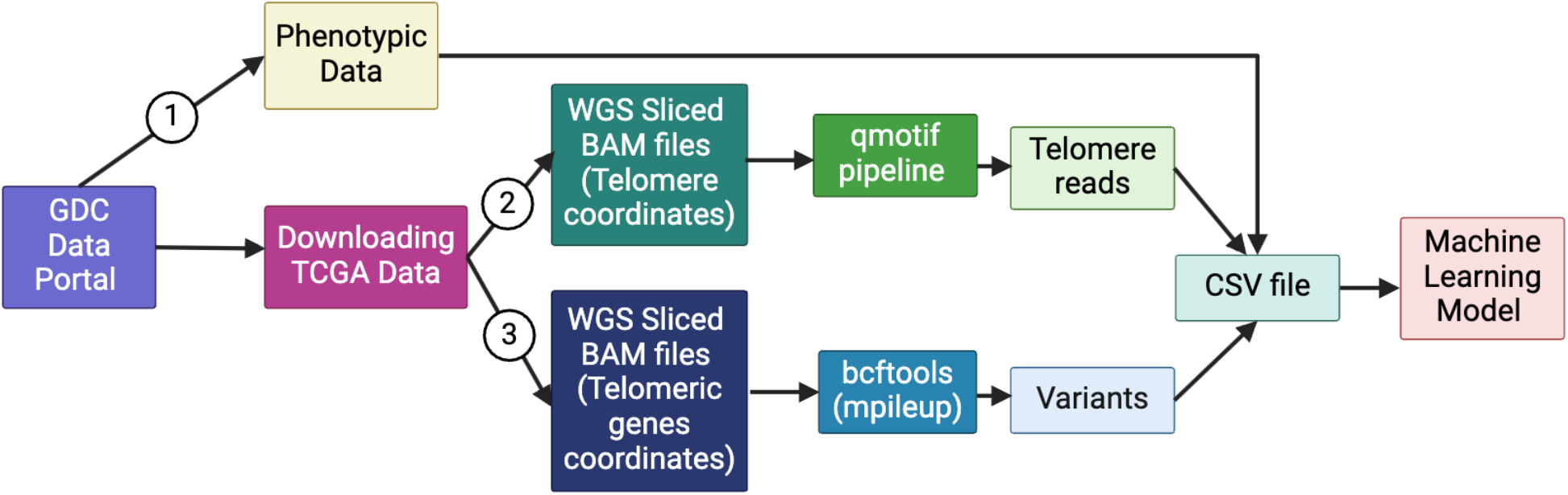
TeloQuest Pipeline – a step-by-step rendition of the analysis pipeline for extracting telomeric read content and genomic variants to train our machine learning model for tumor prediction. [Created with BioRender.com].

**Figure 2.**
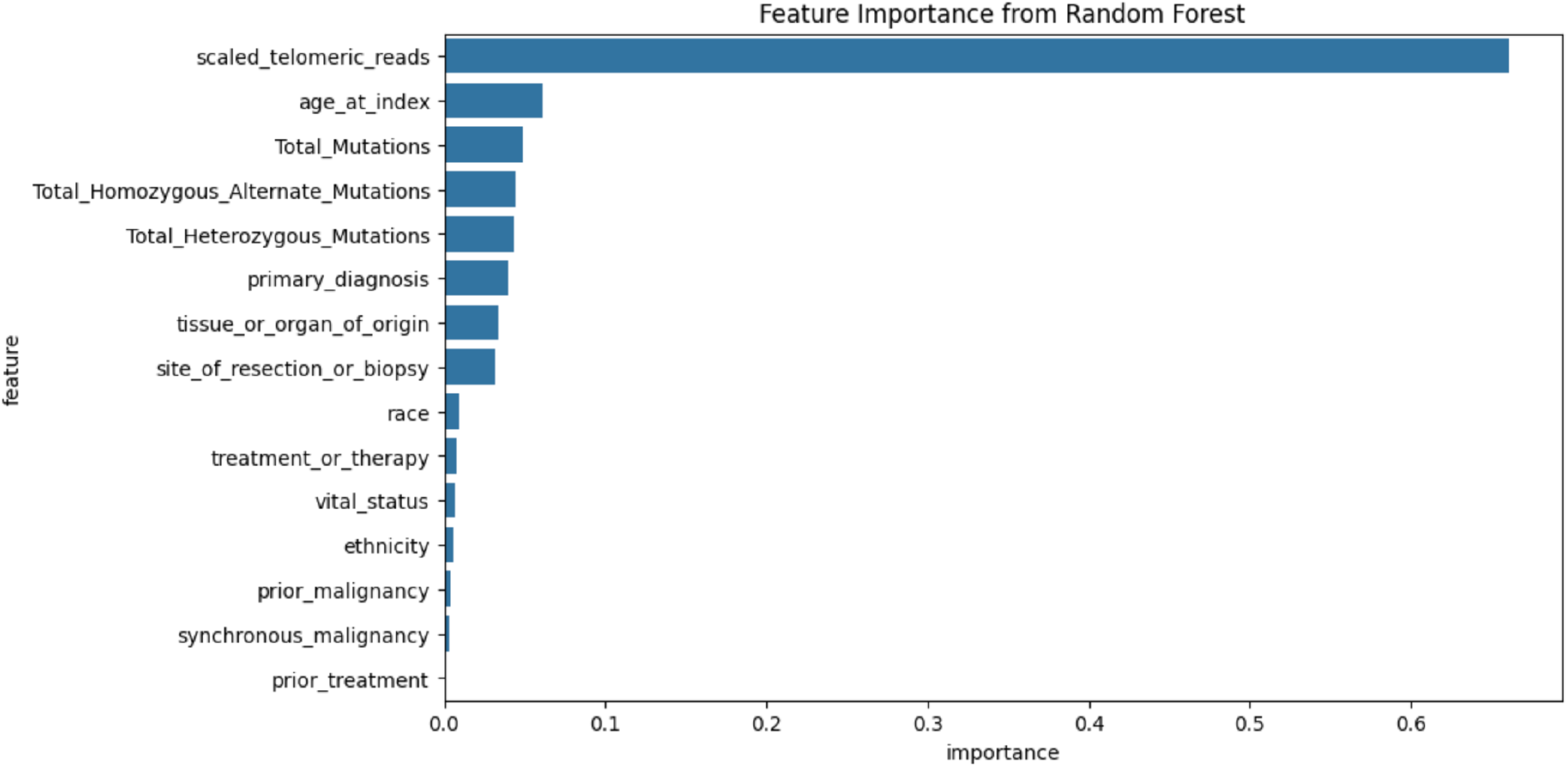
Model attributes that contribute to the greatest feature importance in the Random Forest predictive model of tumor status.

## RESULTS

### Hyperparameter Optimization

The Random Forest classification model was trained to predict binary tumor status (cancer vs. no cancer) using a range of telomeric, genomic, diagnostic, and phenotypic features. The optimal set of hyperparameters included 200 trees (n_estimators), a maximum depth of 10 (max_depth), a feature sampling rate of 50% (max_features = 0.5), a minimum of 2 samples required to split an internal node (min_samples_split = 2), with both bootstrapping (bootstrap = True) and out-of-bag (OOB) error estimation (oob_score = True) enabled. At the optimal classification threshold of 0.4133, our model achieved a test accuracy of 82.62%, a sensitivity (recall) of 79.62%, a specificity of 85.73%, a precision of 85.25%, and an F1 score of 82.34% (Table 3). These results indicate a strong overall predictive performance and a robust ability to detect tumor status while maintaining a high true negative rate.

**Table 3.** Performance Metrics for our Random Forest Classifier at Default, Youden’s Index, Optimal, and High Sensitivity Thresholds.

| Threshold (t) | Sensitivity (Recall) | Specificity (True Negative Rate) | Accuracy | Precision | F1 Score |
| --- | --- | --- | --- | --- | --- |
| Default<br>(t = 0.5) | 72.11% | 92.92% | 82.33% | 91.34% | 80.59% |
| Youden's Index<br>(t = 0.4133) | 79.62% | 85.73% | 82.62% | 85.25% | 82.34% |
| Optimal threshold<br>(t = 0.4133) | 79.62% | 85.73% | 82.62% | 85.25% | 82.34% |
| Enforcing<br>sensitivity of ><br>85%<br>(t = 0.3) | 89.07% | 63.88% | 76.69% | 71.85% | 79.54% |

### Key Features Identified by the Random Forest Model

The five most important features contributing to tumor status prediction were: (1) scaled telomeric read content, (2) age at diagnosis, (3) total number of mutations, (4) total homozygous alternate mutations, and (5) total heterozygous mutations (Figure 2). These variables provided the most predictive power in the model, highlighting the importance of telomeric and mutational data in cancer classification.

### Performance Metrics

The optimal classification threshold, determined using Youden’s Index, was 0.4133. At this threshold, the model achieved a Youden’s Index of 0.6536 (Table 3), reflecting a strong balance between sensitivity and specificity – a critical consideration for minimizing false positives while accurately identifying tumor cases. Additionally, the model’s performance was evaluated using the Area Under the Curve (AUC) from the Receiver Operating Characteristic (ROC) analysis. The AUC was 0.90 (Figure 3), indicating strong discriminative ability and overall model robustness in distinguishing between cancerous and non-cancerous samples.

**Figure 3.**
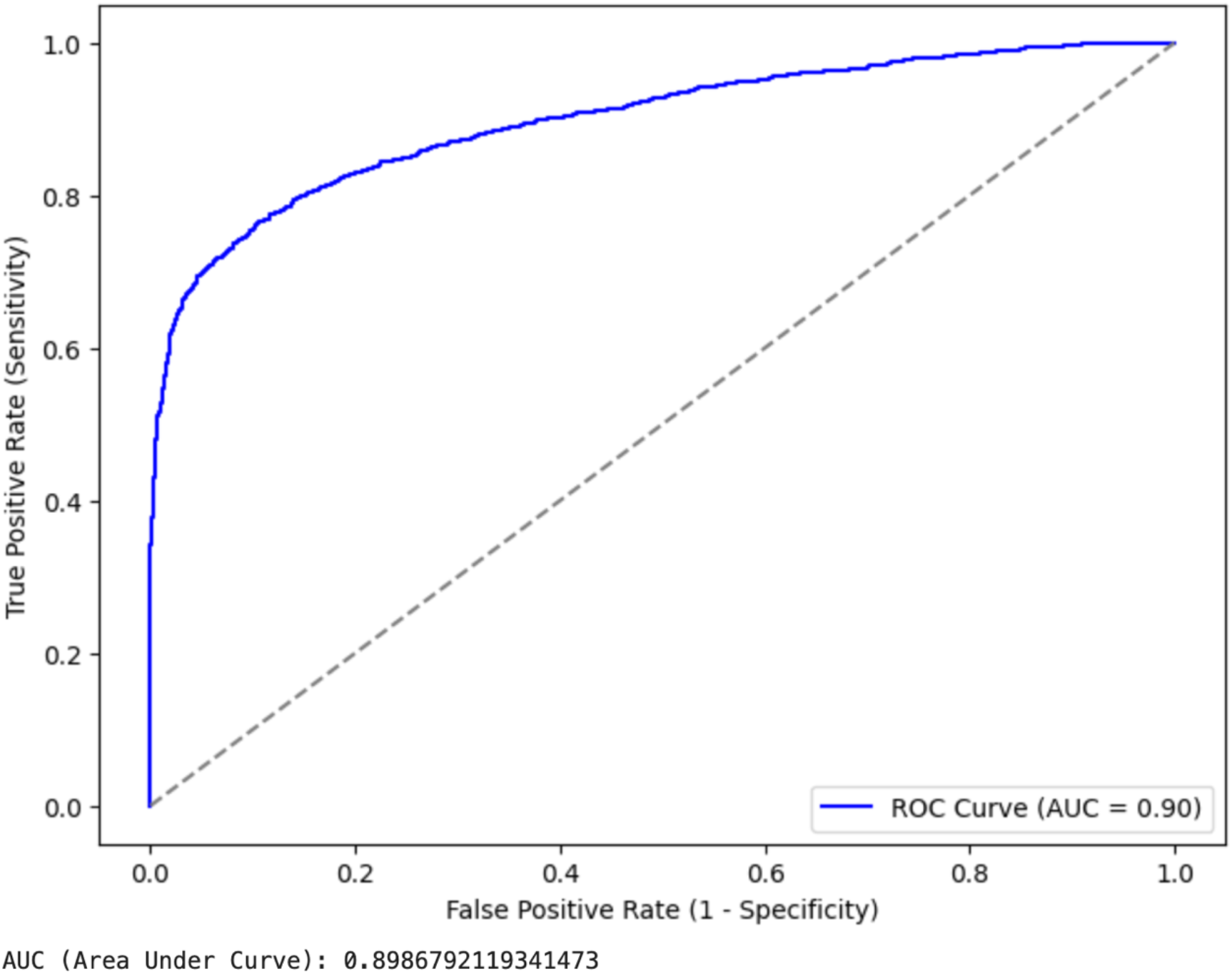
ROC curve (AUC = 0.90) of the Random Forest supervised machine learning model to predict cancer status.

## DISCUSSION

Our primary objective was to develop a model capable of predicting tumor status using telomeric read content, phenotypic data, diagnostic, and genomic information – specifically, variants in telomere length associated genes, from individuals in the TCGA dataset. To achieve this, we compiled a set of 15 features from the GDC Data Portal for each cancer type (Figure 2), including genomic variables such as scaled telomeric length and the total number of homozygous alternate alleles, as well as phenotypic attributes like age at diagnosis. We implemented a Random Forest classification model, chosen for its ability to capture non-linear relationships between predictors and response variables, and for its built-in mechanism to evaluate feature importance. In this framework, feature importance offers a reliable measure of each variable’s contribution to the model’s predictive performance.

In this study, telomere content variation analyses of the TCGA projects showed the following key findings: (1) significant differences in TL between tumor and non-tumor tissues in the same individual, (2) the potential benefits of maintaining a catalog of TLs in cancers and their corresponding normal tissues to improve cancer risk assessment based on TL measurements, and (3) the ability to predict tumor status based on TL variation analysis combined with cancer-related genotypes and phenotypes.

For future investigations, additional attributes, such as socio-economic factors, lifestyle factors, and clinical data, should be explored for incorporation into cancer predictive models. Since machine learning based predictive models can be continually improved, we believe that this straightforward approach has the potential to enhance the communication and interpretation of genetic data in combination with diagnostics. Medical practitioners can use these models to draw inferences about tumor status. At the very least, these predictive models offer a new perspective on predicting cancer. Our Random Forest model demonstrates that genetic summary statistics—including telomere data, total homozygous alternate mutations, and total heterozygous mutations are strong predictors of tumor status and should be included in cancer diagnostics and risk assessments.

The absence of reliable ploidy information is one of the caveats of our current Random Forest model; variant calling at telomere-length associated genes across tumor and non-tumor samples was performed assuming diploidy via bcftools mpileup. Additionally, the Random Forest model here is trained upon 33 different types of cancer, including 10 rare types of cancer. However, we acknowledge that the samples in the TCGA program are primarily derived from the United States population, leading to the underrepresentation of global groups in cancer genomics research. As a result, currently trained models may lack generalizability due to limited genetic diversity in the training data. To ensure that research findings are relevant and beneficial to all populations – not just those already well-represented – there is a critical need to collect more biological samples (e.g., DNA or tumor tissue) from underrepresented groups. We propose that initiatives like TCGA be undertaken globally to capture broader genetic diversity and better understand the complex relationships between genotype and phenotype in cancer.

We show however that a supervised machine learning approach that combines big genomic and phenotypic data can be utilized to develop new preventive strategies and treatment approaches for other telomere-related diseases based on individual genetic telomere content data. Individual telomere content risk could also be valuable for early disease detection beyond cancer, such as in cardiovascular disease and Alzheimer’s disease. Neural networks, especially convolutional neural networks (CNNs), are highly effective for image-related tasks and can be created by incorporating clinical data such as slide images to improve disease diagnostics.

We also acknowledge a tradeoff in the supervised learning model between false positives and false negatives; decreasing one increases the other. However, since this model predicts cancer, we contend that false negative rates are more costly and potentially lethal.

While predictive models, such as the one developed in this study, offer valuable insights and can enhance clinical decision-making, we do not advocate for these models as replacements for medical practitioners. Instead, we envision these predictive machine learning tools as support systems that provide practitioners with additional information to complement their clinical expertise. By integrating data-driven insights with the expert judgment and experience of healthcare providers, we aim to improve patient outcomes while maintaining the central role of human expertise in healthcare.

## ACKNOWLEDGEMENTS

We would like to extend our heartfelt gratitude to the individuals who made this research possible. To the patients who consented to have their samples analyzed by The Cancer Genome Atlas Network, your contributions are the foundation of this research. This work is based on data generated by the TCGA Research Network: https://www.cancer.gov/tcga. We would also like to thank Dr. Kyle Hasenstab for his valuable recommendations on earlier versions of this manuscript.

## AUTHOR CONTRIBUTIONS

AS conceived the study, obtained funding, directed research, and edited the manuscript. PS performed all the research, designed and implemented the software pipelines, collated results, and wrote the manuscript.

## DATA AVAILABILITY

The machine learning pipeline developed to predict tumor status by analyzing telomere content variation can be accessed at (https://github.com/paribytes/TeloQuest). Whole genomic data used in this project can be requested from GDC Data Portal (https://portal.gdc.cancer.gov/).

## REFERENCES

Aunan, J. R., Watson, M. M., Hagland, H. R., & Søreide, K. (2016). Molecular and biological hallmarks of ageing. British Journal of Surgery, 103(2), e29–e46. 10.1002/bjs.10053

Aviv, A., Anderson, J. J., & Shay, J. W. (2017). Mutations, Cancer and the Telomere Length Paradox. Trends in Cancer, 3(4), 253–258. 10.1016/j.trecan.2017.02.005

Burren, O. S., Dhindsa, R. S., Deevi, S. V. V., Wen, S., Nag, A., Mitchell, J., Hu, F., Loesch, D. P., Smith, K. R., Razdan, N., Olsson, H., Platt, A., Vitsios, D., Wu, Q., AstraZeneca Genomics Initiative, Ågren, R., Anderson-Dring, L., Atanur, S., Baker, D., … Petrovski, S. (2024). Genetic architecture of telomere length in 462,666 UK Biobank whole-genome sequences. Nature Genetics, 56(9), 1832–1840. 10.1038/s41588-024-01884-7

Danecek, P., Bonfield, J. K., Liddle, J., Marshall, J., Ohan, V., Pollard, M. O., Whitwham, A., Keane, T., McCarthy, S. A., Davies, R. M., & Li, H. (2021). Twelve years of SAMtools and BCFtools. GigaScience, 10(2), giab008. 10.1093/gigascience/giab008

Factor-Litvak, P., Susser, E., Kezios, K., McKeague, I., Kark, J. D., Hoffman, M., Kimura, M., Wapner, R., & Aviv, A. (2016). Leukocyte Telomere Length in Newborns: Implications for the Role of Telomeres in Human Disease. Pediatrics, 137(4), e20153927. 10.1542/peds.2015-3927

Gomes, N. M. V., Ryder, O. A., Houck, M. L., Charter, S. J., Walker, W., Forsyth, N. R., Austad, S. N., Venditti, C., Pagel, M., Shay, J. W., & Wright, W. E. (2011). Comparative biology of mammalian telomeres: Hypotheses on ancestral states and the roles of telomeres in longevity determination. Aging Cell, 10(5), 761–768. 10.1111/j.1474-9726.2011.00718.x

Greaves, M. (2007). Darwinian medicine: A case for cancer. Nature Reviews Cancer, 7(3), 213–221. 10.1038/nrc2071

Hansen, M. E. B., Hunt, S. C., Stone, R. C., Horvath, K., Herbig, U., Ranciaro, A., Hirbo, J., Beggs, W., Reiner, A. P., Wilson, J. G., Kimura, M., De Vivo, I., Chen, M. M., Kark, J. D., Levy, D., Nyambo, T., Tishkoff, S. A., & Aviv, A. (2016). Shorter telomere length in Europeans than in Africans due to polygenetic adaptation. Human Molecular Genetics, 25(11), 2324–2330. 10.1093/hmg/ddw070

Hedrick, P. W. (2006). Genetic Polymorphism in Heterogeneous Environments: The Age of Genomics. Annual Review of Ecology, Evolution, and Systematics, 37(1), 67–93. 10.1146/annurev.ecolsys.37.091305.110132

Hinrichs, A. S. (2006). The UCSC Genome Browser Database: Update 2006. Nucleic Acids Research, 34(90001), D590–D598. 10.1093/nar/gkj144

Holmes, O., Nones, K., Tang, Y. H., Loffler, K. A., Lee, M., Patch, A.-M., Dagg, R. A., Lau, L. M. S., Leonard, C., Wood, S., Xu, Q., Pickett, H. A., Reddel, R. R., Barbour, A. P., Grimmond, S. M., Waddell, N., & Pearson, J. V. (2022). qmotif: Determination of telomere content from whole-genome sequence data. Bioinformatics Advances, 2(1), vbac005. 10.1093/bioadv/vbac005

Kidd, J. M., Gravel, S., Byrnes, J., Moreno-Estrada, A., Musharoff, S., Bryc, K., Degenhardt, J. D., Brisbin, A., Sheth, V., Chen, R., McLaughlin, S. F., Peckham, H. E., Omberg, L., Bormann Chung, C. A., Stanley, S., Pearlstein, K., Levandowsky, E., Acevedo-Acevedo, S., Auton, A., … Bustamante, C. D. (2012). Population Genetic Inference from Personal Genome Data: Impact of Ancestry and Admixture on Human Genomic Variation. The American Journal of Human Genetics, 91(4), 660–671. 10.1016/j.ajhg.2012.08.025

Kroupa, M., Rachakonda, S. K., Liska, V., Srinivas, N., Urbanova, M., Jiraskova, K., Schneiderova, M., Vycital, O., Vymetalkova, V., Vodickova, L., Kumar, R., & Vodicka, P. (2019). Relationship of telomere length in colorectal cancer patients with cancer phenotype and patient prognosis. British Journal of Cancer, 121(4), 344–350. 10.1038/s41416-019-0525-3

Maciejowski, J., & De Lange, T. (2017). Telomeres in cancer: Tumour suppression and genome instability. Nature Reviews Molecular Cell Biology, 18(3), 175–186. 10.1038/nrm.2016.171

Mangino, M., Christiansen, L., Stone, R., Hunt, S. C., Horvath, K., Eisenberg, D. T. A., Kimura, M., Petersen, I., Kark, J. D., Herbig, U., Reiner, A. P., Benetos, A., Codd, V., Nyholt, D. R., Sinnreich, R., Christensen, K., Nassar, H., Hwang, S.-J., Levy, D., … Aviv, A. (2015). DCAF4, a novel gene associated with leucocyte telomere length. Journal of Medical Genetics, 52(3), 157–162. 10.1136/jmedgenet-2014-102681

McGrath, M., Wong, J. Y. Y., Michaud, D., Hunter, D. J., & De Vivo, I. (2007). Telomere Length, Cigarette Smoking, and Bladder Cancer Risk in Men and Women. Cancer Epidemiology, Biomarkers & Prevention, 16(4), 815–819. 10.1158/1055-9965.epi-06-0961

McNally, E. J., Luncsford, P. J., & Armanios, M. (2019). Long telomeres and cancer risk: The price of cellular immortality. Journal of Clinical Investigation, 129(9), 3474–3481. 10.1172/JCI120851

McQuillan, M. A., Verhulst, S., Hansen, M. E. B., Beggs, W., Meskel, D. W., Belay, G., Nyambo, T., Mpoloka, S. W., Mokone, G. G., Fokunang, C., Njamnshi, A. K., Chanock, S. J., Aviv, A., & Tishkoff, S. A. (2024). Association between telomere length and Plasmodium falciparum malaria endemicity in sub-Saharan Africans. The American Journal of Human Genetics, 111(5), 927–938. 10.1016/j.ajhg.2024.04.003

Okamoto, K., & Seimiya, H. (2019). Revisiting Telomere Shortening in Cancer. Cells, 8(2), 107. 10.3390/cells8020107

Rachakonda, S., Kong, H., Srinivas, N., Garcia-Casado, Z., Requena, C., Fallah, M., Heidenreich, B., Planelles, D., Traves, V., Schadendorf, D., Nagore, E., & Kumar, R. (2018). Telomere length, telomerase reverse transcriptase promoter mutations, and melanoma risk. Genes, Chromosomes and Cancer, 57(11), 564–572. 10.1002/gcc.22669

Stanley, S. E., & Armanios, M. (2015). The short and long telomere syndromes: Paired paradigms for molecular medicine. Current Opinion in Genetics & Development, 33, 1–9. 10.1016/j.gde.2015.06.004

Stranger, B. E., Montgomery, S. B., Dimas, A. S., Parts, L., Stegle, O., Ingle, C. E., Sekowska, M., Smith, G. D., Evans, D., Gutierrez-Arcelus, M., Price, A., Raj, T., Nisbett, J., Nica, A. C., Beazley, C., Durbin, R., Deloukas, P., & Dermitzakis, E. T. (2012). Patterns of Cis Regulatory Variation in Diverse Human Populations. PLoS Genetics, 8(4), e1002639. 10.1371/journal.pgen.1002639

The Telomeres Mendelian Randomization Collaboration, Haycock, P. C., Burgess, S., Nounu, A., Zheng, J., Okoli, G. N., Bowden, J., Wade, K. H., Timpson, N. J., Evans, D. M., Willeit, P., Aviv, A., Gaunt, T. R., Hemani, G., Mangino, M., Ellis, H. P., Kurian, K. M., Pooley, K. A., Eeles, R. A., … Davey Smith, G. (2017). Association Between Telomere Length and Risk of Cancer and Non-Neoplastic Diseases: A Mendelian Randomization Study. JAMA Oncology, 3(5), 636. 10.1001/jamaoncol.2016.5945

Tomczak, K., Czerwińska, P., & Wiznerowicz, M. (2015). Review The Cancer Genome Atlas (TCGA): An immeasurable source of knowledge. Współczesna Onkologia, 1A, 68–77. 10.5114/wo.2014.47136

Verhulst, S., Susser, E., Factor-Litvak, P. R., Simons, M. J., Benetos, A., Steenstrup, T., Kark, J. D., & Aviv, A. (2015). Commentary: The reliability of telomere length measurements. International Journal of Epidemiology, 44(5), 1683–1686. 10.1093/ije/dyv166

Victorelli, S., & Passos, J. F. (2017). Telomeres and Cell Senescence—Size Matters Not. EBioMedicine, 21, 14–20. 10.1016/j.ebiom.2017.03.027

Xu, L., Li, S., & Stohr, B. A. (2013). The Role of Telomere Biology in Cancer. Annual Review of Pathology: Mechanisms of Disease, 8(1), 49–78. 10.1146/annurev-pathol-020712-164030

Zhu, X., Han, W., Xue, W., Zou, Y., Xie, C., Du, J., & Jin, G. (2016). The association between telomere length and cancer risk in population studies. Scientific Reports, 6(1). 10.1038/srep22243

