## Supplementary Data for "A NOVEL MACHINE LEARNING APPROACH FOR TUMOR DETECTION BASED ON TELOMERIC SIGNATURES"

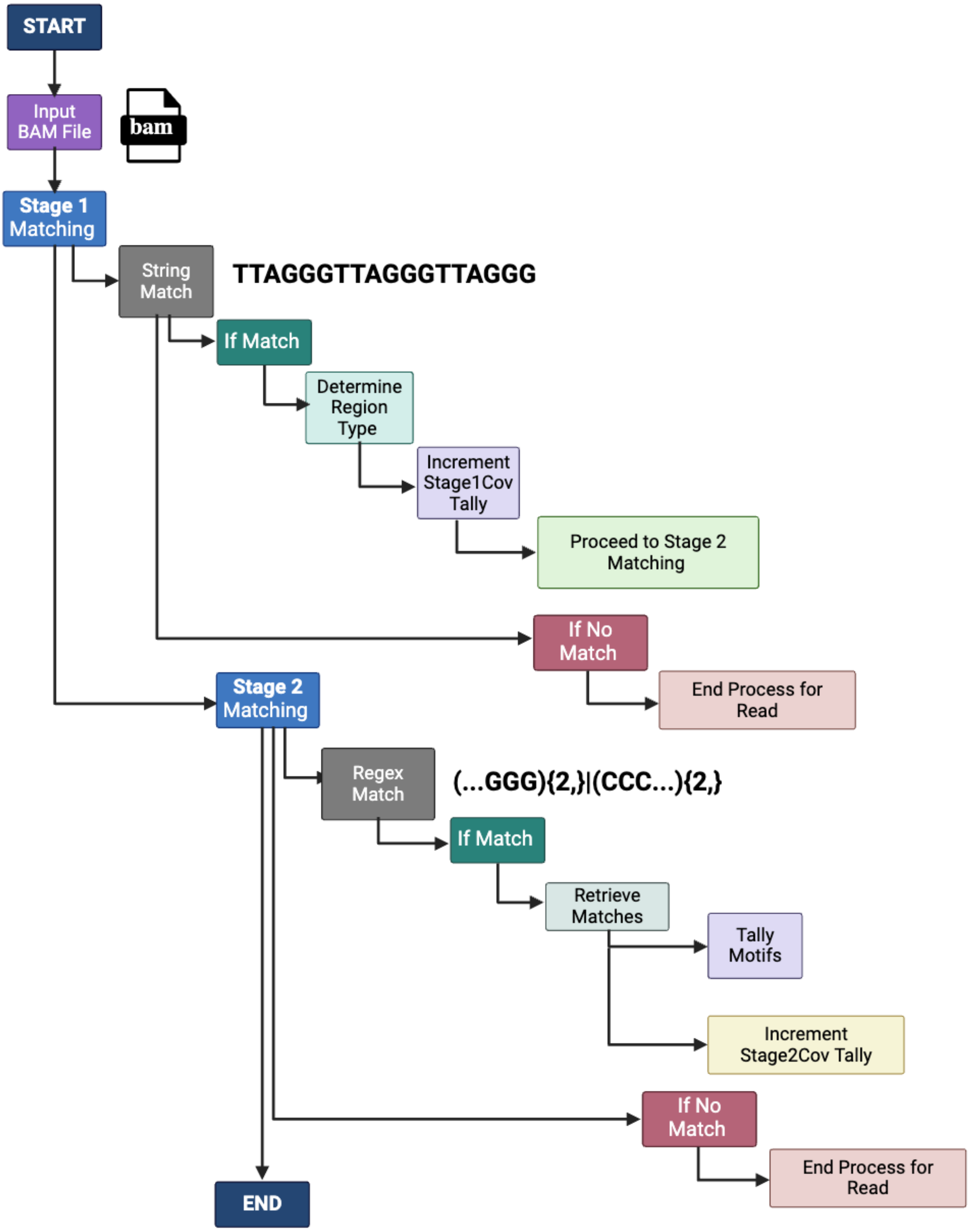


**Supplementary Figure S1.** Workflow of the telomere content variation pipeline, that incorporates qmotif to estimate variation from TCGA data [Created with BioRender.com].

**Supplementary Table S1.** Summary of phenotypic predictor variables included in the Random Forest classifier (Source: https://docs.cancergenomicscloud.org/docs/tcga-grch38-metadata)

| Feature | Entity | Description |
|---|---|---|
| Age at diagnosis | Diagnosis | The age in years of the case at the initial pathological diagnosis of the disease or cancer. |
| Ethnicity | Demographic | A socially-defined category of people based on common ancestral, cultural, biological, and social factors. |
| Gender | Demographic | The collection of behaviors and attitudes that distinguish people on the basis of the societal roles expected for the two sexes. |
| Race | Demographic | A classification of humans characterized by certain heritable traits, common history, nationality, or geographic distribution. |
| Vital status | Diagnosis | The state of being living or deceased for Cases that are part of the investigation. |
| Primary diagnosis | Diagnosis | Text term for the structural pattern of cancer cells used to define a microscopic diagnosis. |
| Prior malignancy | Diagnosis | Text term to describe the patient's history of prior cancer diagnosis and the spatial location of any previous cancer occurrence. |
| Prior treatment | Diagnosis | A yes/no/unknown/not applicable indicator related to if the patients were given prior treatment. |
| Site of resection or biopsy | Diagnosis | The topography code which describes the anatomical site of origin of the neoplasm according to the third edition of the International Classification of Diseases for Oncology. |
| Synchronous malignancy | Diagnosis | A yes/no/unknown indicator if a patient has more than one primary malignancy diagnosed within six months of each other. |

| | | |
|---|---|---|
| Tissue or organ of origin | Diagnosis | The text term that describes the anatomic site of the tumor or disease. |
| Treatment or therapy | Treatment | A yes/no/unknown/not applicable indicator related to the administration of therapeutic agents received before the body specimen was collected. |
| Treatment type | Treatment | Pharmaceutical therapy or Radiation therapy given |
| Tumor status | Diagnosis | The condition or state of the tumor at a particular time. |

**Supplementary Table S2.** Gene symbols, names and genomic coordinates of the 15 telomere related genes for which variants have been captured from the paper by Burren et al. (2024).

| No. | Gene Symbol | Gene Full Name | Genomic coordinates |
|---|---|---|---|
| 1. | DCLRE1B | DNA cross-link repair 1B | chr1:113905326-113914086 |
| 2. | NAF1 | Nuclear assembly factor 1 ribonucleoprotein | chr4:163128669-163166890 |
| 3. | TERT | Telomerase reverse transcriptase | chr5:1253167-1295068 |
| 4. | G3BP1 | G3BP stress granule assembly factor 1 | chr5:151771954-151812785 |
| 5. | ZNF451 | Zinc finger protein 451 | chr6:57090188-57170305 |
| 6. | POT1 | Protection of telomeres 1 | chr7:124822386-124929825 |
| 7. | TERF1 | Telomeric repeat binding factor 1 | chr8:73008864-73048123 |
| 8. | STN1 | STN1 subunit of CST complex | chr10:103877569-103918184 |
| 9. | ATM | ATM serine/threonine kinase | chr11:108223067-108369102 |
| 10. | TINF2 | TERF1 interacting nuclear factor 2 | chr14:24239643-24242623 |

| | | | |
|---|---|---|---|
| 11. | PARN | Poly(A)-specific ribonuclease | chr16:14435701-14630260 |
| 12. | CTC1 | CST telomere replication complex component 1 | chr17:8224815-8248056 |
| 13. | BRIP1 | BRCA1 interacting DNA helicase 1 | chr17:61679139-61863528 |
| 14. | SAMHD1 | SAM and HD domain containing deoxynucleoside triphosphate triphosphohydrolase 1 | chr20:36890229-36951708 |
| 15. | RTEL1 | Regulator of telomere elongation helicase 1 | chr20:63658312-63696245 |

**Supplementary Table S3.** Whole genomic data and project type of each cancer available in TCGA program analyzed for telomere content variation and variants.

| Number | Cancer Type (Project code) | Primary Site | Total no. of cases | Total no. of BAM files in GDC data portal | Total BAM files analyzed (Control and tumor samples) |
|---|---|---|---|---|---|
| 1. | TCGA – ACC | Adrenal gland | 74 | 148 | 148 |
| 2. | TCGA – BLCA | Bladder | 411 | 1,695 | 820 |
| 3. | TCGA – BRCA | Breast | 952 | 2,105 | 1,904 |
| 4. | TCGA – CESC | Cervix uteri | 271 | 654 | 542 |
| 5. | TCGA – CHOL | Gall bladder | 46 | 92 | 92 |
| 6. | TCGA – COAD | Colon | 370 | 935 | 740 |
| 7. | TCGA – DLBC | Lymphoid | 42 | 98 | 84 |
| 8. | TCGA – ESCA | Esophagus | 118 | 281 | 236 |
| 9. | TCGA – GBM | Brain | 344 | 798 | 688 |

| | | | | | |
|---|---|---|---|---|---|
| 10. | TCGA – HNSC | Mouth | 482 | 1,221 | 964 |
| 11. | TCGA – KICH | Kidney | 86 | 270 | 172 |
| 12. | TCGA – KIRC | Kidney | 124 | 260 | 248 |
| 13. | TCGA – KIRP | Kidney | 216 | 489 | 432 |
| 14. | TCGA – LAML | Bone marrow | 50 | 118 | 100 |
| 15. | TCGA – LGG | Brain | 461 | 1,100 | 920 |
| 16. | TCGA – LIHC | Liver | 324 | 722 | 648 |
| 17. | TCGA – LUAD | Lung | 464 | 1,186 | 928 |
| 18. | TCGA – LUSC | Lung | 337 | 706 | 674 |
| 19. | TCGA – MESO | Heart | 73 | 146 | 146 |
| 20. | TCGA – OV | Ovary | 362 | 787 | 724 |
| 21. | TCGA – PAAD | Pancreas | 173 | 429 | 346 |
| 22. | TCGA – PCPG | Multiple sites | 165 | 330 | 330 |
| 23. | TCGA – PRAD | Prostate | 414 | 1056 | 828 |
| 24. | TCGA – READ | Colon | 143 | 376 | 284 |
| 25. | TCGA – SARC | Multiple sites | 223 | 514 | 446 |
| 26. | TCGA – SKCM | Skin | 223 | 524 | 446 |
| 27. | TCGA – STAD | Stomach | 436 | 1,446 | 872 |
| 28. | TCGA – TGCT | Testis | 237 | 478 | 248 |

| | | | | | |
|---|---|---|---|---|---|
| 29. | TCGA – THCA | Thyroid | 477 | 1,249 | 954 |
| 30. | TCGA – THYM | Thymus | 111 | 222 | 222 |
| 31. | TCGA – UCEC | Corpus uteri | 482 | 1,225 | 964 |
| 32. | TCGA – UCS | Uterus | 50 | 100 | 100 |
| 33. | TCGA – UVM | Eye | 75 | 244 | 150 |
| Total | | | 8,816 | 22,004 | 17,400 |
